# Non-enzymatic roles of human RAD51 at stalled replication forks

**DOI:** 10.1101/359380

**Authors:** Jennifer M. Mason, Yuen-Ling Chan, Ralph W. Weichselbaum, Douglas K. Bishop

## Abstract

The central recombination enzyme RAD51 has been implicated in replication fork processing and restart in response to replication stress. Here, we use a separation-of-function allele of RAD51 that retains DNA binding, but not strand exchange activity, to reveal mechanistic aspects of RAD51’s roles in the response to replication stress. We find that cells lacking RAD51 strand exchange activity protect replication forks from MRE11-dependent degradation, as expected from previous studies. Unexpectedly we find that RAD51’s strand exchange activity is not required to convert stalled forks to a form that can be degraded by DNA2. Such conversion was shown previously to require replication fork reversal, supporting a model in which fork reversal depends on a non-enzymatic function of RAD51. We also show RAD51 promotes replication restart by both strand exchange-dependent and strand exchange-independent mechanisms.

## INTRODUCTION

The complete and accurate replication of the genome is essential to maintain genome integrity. Replication forks face many obstacles that can result in replication fork stalling or replication fork collapse. Proteins initially identified on the basis of their roles in homologous recombination (HR) are now known to have key functions during replication stress^1,2^. HR proteins act to protect and remodel stalled replication forks, and re-construct functional replication forks following fork collapse. As a result of these activities, HR proteins are critical to the ability of cells to restart stalled and collapsed replication forks.

The central HR protein, RAD51, forms helical nucleoprotein filaments on tracts of single strand DNA (ssDNA), such as those formed by nucleolytic processing of the DNA ends at the sites of DNA double strand breaks (DSBs). Once RAD51 filaments form on tracts of ssDNA, the protein alters the structure of the ssDNA, allowing the nucleoprotein filament to catalyze a homology search to identify an identical or nearly-identical sequence in duplex DNA, and then carry out exchange of the bound ssDNA strand with the “like” strand of the homologous duplex^3^. In this way, the homology search and strand exchange activity of RAD51 acts to form a homologous joint between a broken chromatid and its intact sister chromatid, leading to accurate repair of the DSB.

In addition to its role in repair of DSBs, RAD51 has three separate roles at stalled replication forks. First, RAD51 promotes replication fork reversal^4,5^. Replication fork reversal involves branch migration in the direction opposite to replication forming a Holliday junction-containing “chicken foot” structure. Second, RAD51 protects tracts of newly-synthesized “nascent” DNA from degradation. Initial resection of the reversed forks by MRE11, EXO1, and DNA2 produces a ssDNA overhang at reversed forks^5,6^. The formation of a stable RAD51 filament on the resulting ssDNA overhang prevents extensive degradation of reversed forks. Nascent ssDNA degradation occurs in cells with partial inhibition of RAD51 expression or activity in response a variety of DNA damage agents^7-11,16^. Nascent DNA degradation also occurs in cells lacking proteins required to load RAD51 on ssDNA such as BRCA1,BRCA2, FANCD2, and the RAD51 mediators including RAD51C and XRCC2^5,7,12-15^. The degradation phenotype observed in these cells results from the inefficient nucleation and/or stabilization of RAD51 filaments at the reversed fork^5,14,15,17^. Three nucleases involved in DNA end resection, MRE11, EXO1, and DNA2 are responsible for the degradation of nascent DNA strands. MRE11 and EXO1 degrade forks in cells with defects in RAD51 filament formation and stabilization. DNA2 degradation has been observed under two conditions. First, mutant cells in which RAD51 is efficiently loaded onto reversed forks, but filament stability is decreased results in DNA2-dependent degradation^18^. Second, prolonged HU treatment results in DNA2-dependent degradation that occurs even without manipulations that decrease RAD51 filament stability^6,11^. Depletion of RADX, a single stranded binding protein that negatively regulates RAD51 at reversed replication forks, prevents excessive MRE11- and DNA2-dependent degradation that results from defects that reduce RAD51 filament formation or RAD51 filament stability^11^. However, depletion of RADX did not prevent DNA2-dependent degradation after prolonged HU treatment, indicating degradation occurs independently of RAD51 filament stability. Finally, RAD51 plays a role in restart of stalled or collapsed replication forks^19^. Collapse of replication forks may sometimes occur as a consequence of collision of forks with a single strand nick in the template and has been to shown to occur as a consequence of enzymatic cleavage of reversed forks by structure-specific endonucleases including MUS81^20-22^. Repair of the resulting DSB by HR to reinstate the replication fork requires RAD51 and MRE11 activity^19,23,24^.

Although it is clear that RAD51 is required for protection of nascent DNA strands from MRE11, replication fork remodeling, and fork restart, the molecular mechanisms underlying RAD51’s role in each of these functions remains to be determined. One particularly important question is whether and to what degree RAD51’s homology search and strand exchange activity is required for each its roles during the replication stress response. The ability of RAD51 to promote nuclease protection at stalled forks and/or HR can be separated from its ability to promote fork reversal. Partial knockdown of RAD51 resulted in MRE11-dependent degradation of replication forks, but knockdown to low levels of RAD51 rescues fork deprotection by preventing replication fork reversal^11^. RAD51-dependent replication fork reversal is BRCA2-independent^5,17^ even though protection of reversed forks from pathological degradation by MRE11 is BRCA2-dependent. In addition, replication-associated sister chromatid exchange has been observed in BRCA2-deficient cells that display defects in DSB-dependent HR and RAD51 immunostaining focus formation^25^. Another important study showed cells that co-express RAD51 WT with RAD51 T131P, a dominant-negative allele of RAD51 that forms unstable nucleoprotein filaments due to constitutive ATPase activity, are HR proficient, undergo fork reversal, but do not protect forks from degradation by MRE11 or DNA2^5,11,18^. However, the ability of RAD51 T131P to specifically disrupt fork protection is only seen in cells co-expressing RAD51 WT at a ratio that delays RAD51 focus formation, but does not inhibit HR (e.g. in heterozygous, patient-derived cells)^18^; RAD51 T131P protein does not form immunostaining foci and high levels of mutant protein inhibits focus formation of endogenous RAD51. This finding suggests that RAD51 T131P partially inhibits the function of wildtype RAD51 resulting in the observed separation of function. Another mutant allele of RAD51, RAD51 K133R, which is defective in ATP hydrolysis was found to rescue MRE11-dependent fork protection in FANCD2-deficient and BRCA2-deficient cells. However, human RAD51 K133R retains strand exchange activity *in vitro*^26,27^ and can promote significant amounts of HR *in vivo,* although human RAD51 K133R-mediated HR has only been observed in chicken DT40 cells^27^. Given that RAD51 K133R lacks ATPase activity which is involved in promoting filament disassembly^26^, it is possible that the HR defect observed in human cells expressing the mutant protein results from a post-strand exchange block to RAD51 filament disassembly rather than from a defect in strand exchange. Thus, the prior work that separated functions of RAD51 during replication stress did not specifically test the role of the protein’s enzymatic homology search and strand exchange activity. In this context it should be noted that both ensemble and single molecule biochemistry have shown that only 8 bp of homology are required for meta-stable homology-recognition by RAD51 family strand exchange proteins^28,29^. Given the binding stoichiometry of RAD51 to DNA (1 protomer per 3 nucleotides), filaments too short to be detected as immunostaining foci could, in principle, promote strand exchange at stalled replication forks. Thus, strand exchange could be a residual activity of partially-defective RAD51 during replication stress, even in cells defective in RAD51 focus formation, such as BRCA2-deficient cells.

During strand exchange reaction, RAD51 utilizes two distinct binding sites. RAD51 forms nucleoprotein filaments on resected single strand DNA via a high-affinity binding site (site I). The RAD51 nucleoprotein filament searches intact dsDNA for homologous sequences by binding a second low affinity binding site (site II), consisting of a cluster of positively charged residues the lie on the interior surface of the helical filament^30^. This cluster binds backbone phosphates of dsDNA regions being searched, and also continues to bind the out-going ssDNA strand once strand exchange occurs^31^. *In S. cerevisiae*, RAD51 containing mutations in site II (II3A) retains the ability to form nucleoprotein filaments, but is fully-defective in homology search and strand exchange *in vitro* and highly-defective in mitotic HR *in vivo*^30^.

Here, we disrupt the secondary binding site on human RAD51 to determine the role of strand exchange activity in response to replication stress. Our results provide definitive evidence that the strand exchange activity of RAD51 is not required to protect stalled forks from MRE11 degradation, as expected. Surprisingly, we also show that strand exchange activity is not required for RAD51-dependent remodeling of stalled forks to a form that permits DNA2-dependent degradation of nascent DNA. In contrast, we provide evidence that RAD51 promotes replication restart by both strand exchange-dependent and strand exchange-independent mechanisms. Cytological and knock-down experiments show that cells expressing a strand exchange-defective form of RAD51 accumulate collapsed replication forks and undergo frequent new origin firing in response to HU-induced replication stress.

## RESULTS

### hsRAD51-II3A retains significant DNA binding activity, but is defective for D-loop formation

To characterize the molecular functions of human RAD51’s DNA binding and strand exchange activities during replication stress, we constructed an allele of human RAD51 corresponding to the *S. cerevisiae rad51-II3A* allele^30^. The corresponding human RAD51-II3A (hsRAD51-II3A) protein has 3 amino acid residues, R130, R303, and K313 changed to alanines. To determine if the mutant human protein (hsRAD51-II3A) had the same properties as its budding yeast counterpart, we purified hsRAD51-WT and hsRAD51-II3A proteins (Supplemental Figure 1a). We analyzed the nucleoprotein filament forming and strand exchange activities of the two forms of RAD51 using biochemical assays. Using fluorescence polarization (FP), we measured binding of the two forms of the protein to a Alexa84-tagged 88-nt ssDNA oligo^32^. Titration showed that the two proteins have similar binding activities to this oligo, the apparent Kd’s for hsRAD51-WT and hsRAD51-II3A were 57 ± 1 nM and 132 ± 5 nM, respectively (Figure 1a). Thus, hsRAD51-II3A displays only a modest DNA binding defect in this assay. Next, we examined the homology search and strand exchange activity of hsRAD51-II3A with a D-loop assay that employs a 90-nt single strand oligonucleotide and a 4.4-kb supercoiled plasmid carrying a dsDNA sequence identical to the sequence of the oligonucleotide (Figure 1b). In this assay, hsRAD51-II3A exhibited 840-fold less D-loop activity compared to hsRAD51-WT (0.06% vs. 17% of plasmid DNA forming D-loops, respectively). Together the results demonstrate that, like its budding yeast counterpart, hsRAD51-II3A retains DNA binding, but not strand exchange activity *in vitro*.

**Figure 1.**
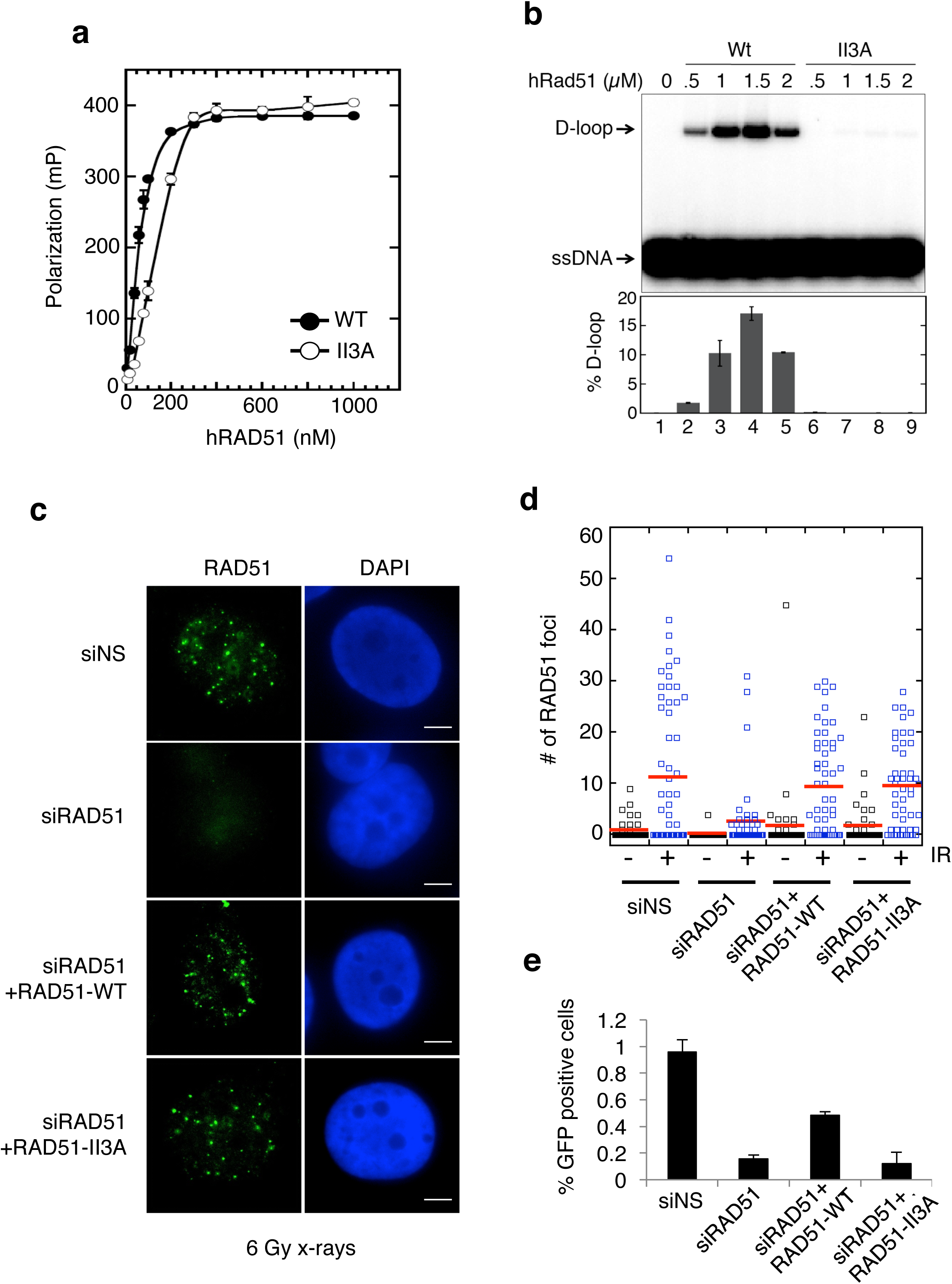
hsRAD51-II3A retains ssDNA binding activity, but is defective for HR. (a) Human RAD51 binding to ssDNA was determined by fluorescence polarization. Protein at various concentrations was incubated with fluorescein-tagged 84-mer ssDNA (200 nM nucleotides or 2.4 nM molecules) in buffer B and incubated at 37°C for 30 min. Error bars represent standard deviation from triplicate experiments. The apparent Kd for hsRAD51-WT is 57 ± 1 nM, for hsRAD51-II3A is 132 ± 5 nM. (b) The D-loop activity of hsRAD51-WT and hsRAD51-II3A was measured in the presence of increasing concentrations of protein. Upper panel: autoradiogram following electrophoretic separation of D-loops from free ^32^P-labelled ssDNA oligo substrate. Lower panel: quantitation of the autoradiogram shown in the upper panel. Error bars represent standard deviation from triplicate experiments. (c) Images depict immuno-staining RAD51 in cells prepared for staining 8 hours after a dose of 6 GY x-rays in the indicated samples. Green-RAD51 Blue-DNA. Scale bar= 5 µm (d) Quantitation of RAD51 foci after the indicated treatments. Red line represents the mean. (e) DR-GFP assay. The percentage of GFP positive cells in each sample are graphed. Error bars represent SEM.

Next, we asked if hsRAD51-II3A displays separation of RAD51’s DNA binding and HR functions *in vivo.* hsRAD51-WT and hsRAD51-II3A were expressed in U2OS cells and the expression of the endogenous RAD51 protein was repressed via siRNA targeting of the 3’UTR of the RAD51 mRNA. A second positive control was carried out by transfection of non-silencing siRNA (siNS). Treatment with RAD51 siRNA reduced expression of the endogenous protein to less than 7% of that observed in the siNS control (Supplemental Figure 1b). Both hsRAD51-WT and hsRAD51-II3A were expressed at the same level, which was ∼5 fold higher than that seen for the endogenous protein in the non-silencing siRNA (siNS) control (Supplemental Figure 1b).

To determine the extent to which the spontaneous distribution of RAD51 differed in cells transfected with RAD51 expression constructs from that observed with endogenous RAD51, we counted RAD51 foci in unselected nuclei. RAD51 typically forms a small number of nuclear immunostaining foci in the absence of induced damage at sites of spontaneous DNA damage or the sites of non-repair-associated RAD51 oligomers. The number of spontaneous RAD51 foci between cells slightly increased in cells expressing hsRAD51-WT (1.5±6.7 RAD51 foci/cell) or hsRAD51-II3A (1.5±4 RAD51 foci/cell), when compared to siNS (0.7±1.8 foci/cell in siNS cells; Figure 1c,d). In addition, a small subpopulation of hsRAD51-WT (3.4±1.5%) and hsRAD51-II3A (2.3±1.3%) transfected cells contained an average of 44±14 elongated RAD51 fibers with contour lengths of 0.5 to 3 microns long (Supplemental Figure 1c). This type of staining pattern was observed previously as a consequence of high levels of RAD51 overexpression and reflects binding of RAD51 to undamaged DNA^33^. This analysis indicated that the level of expression of RAD51 from transfection of siRNA resistant constructs causes only a slight increase in the frequency of spontaneous foci.

Analysis of damage induced RAD51 foci provided evidence that hsRAD51-II3A retains DNA binding activity *in vivo*. Loading of RAD51 protein filaments on ssDNA tracts on the 3’ ssDNA overhang at resected DSBs *in vivo* is conventionally assayed by measuring the number and of RAD51 foci resulting from treatment of cells with agents that induce DNA breaks, such as x-rays^33-36^. The number of RAD51 foci significantly increased after x-ray treatment of siNS transfected cells (Figure 1c,d; 11±14.4 foci/cell IR *vs* 0.7±1.8 foci/cell). Cells depleted of RAD51 exhibited a 4-fold reduction in IR-induced RAD51 focus formation (2.5±6 foci/cell; *p*<0.005). Cells transfected with hsRAD51-WT (9±9.7 foci/cell) showed the same level of x-ray induced foci as the siNS control (11±14.4 foci/cell). Importantly, the number of IR-induced hsRAD51-II3A (9±8.3) foci did not differ significantly from siNS or hsRAD51-WT controls. Together, these results indicate hsRAD51-II3A retains significant DNA binding activity *in vivo*.

To determine if hsRAD51-II3A is defective in HR, we employed the DR-GFP assay^37^. In this assay, HR generates a functional GFP allele following induction of a chromosomal DNA break in one of the two defective copies of GFP carried by the reporter cell line (Supplemental Figure 1d). Double strand breaks were induced in the U2OS-DR-GFP cells by transfection with a plasmid expressing the I-SceI endonuclease. HR efficiency was then measured by flow cytometry as the frequency of GFP expressing cells. RAD51 depletion reduced HR efficiency 6-fold compared to siNS controls (0.97±0.1% GFP positive cells siNS vs 0.16±0.03% siRAD51; *p*-value<0.05; Figure 1e). Expression of hsRAD51-WT in RAD51 depleted cells increased the HR efficiency by 3-fold compared to siRAD51 cells (0.49±0.03 GFP positive cells vs. 0.16±0.03% siRAD51; *p*-value<0.005). In contrast, hsRAD51-II3A did not increase HR efficiency in RAD51-depleted cells (0.12±0.09 % GFP positive cells vs. 0.16±0.03%; *p*-value=0.6) indicating hsRAD51-II3A is defective for HR-mediated repair of DSBs. Consistent with our biochemical observations, these data indicate human hsRAD51-II3A is able to form RAD51 nucleoprotein filaments with normal efficiency, but is defective for HR *in vivo.*

### hsRAD51-II3A protects nascent DNA strands from MRE11-dependent degradation

We next sought to elucidate the molecular function of RAD51’s strand exchange activity during perturbed replication, using the DNA fiber assay to measure the ability of cells treated with the replication inhibitor HU to protect nascent DNA strands from degradation. Nascent DNA undergoes MRE11-mediated degradation in cells that have defects in RAD51 loading and/or stabilization of RAD51 nucleoprotein filaments^5,7,12,14,15^. MRE11-dependent fork degradation in FANCD2 and BRCA2-deficient cells can be rescued by overexpressing RAD51^7,12^. We determined if overexpression of hsRAD51-II3A rescues fork degradation in U2OS cells depleted of FANCD2 after treatment with 4 mM HU for 5 hours (Figure 2a, Supplemental Figure 2a). Consistent with previous studies, we observed replication tract shortening in FANCD2-depleted cells (0.55±0.15 µm vs 0.74±0.16, *p-*value <0.0001). As expected, treatment with mirin (0.78±0.15 µm, *p*<0.0001) and expression of RAD51-WT (0.79±0.18 µm, *p*<0.001) restored replication tracts to lengths observed without HU treatment^12^. Expression of hsRAD51-II3A also restored replication tract length in FANCD2-depleted cells. We verified this result by expressing hsRAD51-WT and hsRAD51-II3A in a BRCA2-deficient (VU423) fibroblast cell line, which has also been shown to undergo MRE11-dependent degradation upon HU induced stress^38^ (Figure 2b). As expected, HU treatment resulted in shortened replication tracts (0.84±18 µm vs 0.62±23 µm, *p-* value <0.0001) that were rescued by treatment with mirin (0.84±16 µm), by expression of hsRAD51-WT (0.85±0.15 µm) and by expression of RAD51-II3A (0.81±0.19 µm). Taken together, these data indicate expression of hsRAD51-II3A rescues MRE11-dependent degradation in HR mutants with destabilized RAD51 filaments.

**Figure 2.**
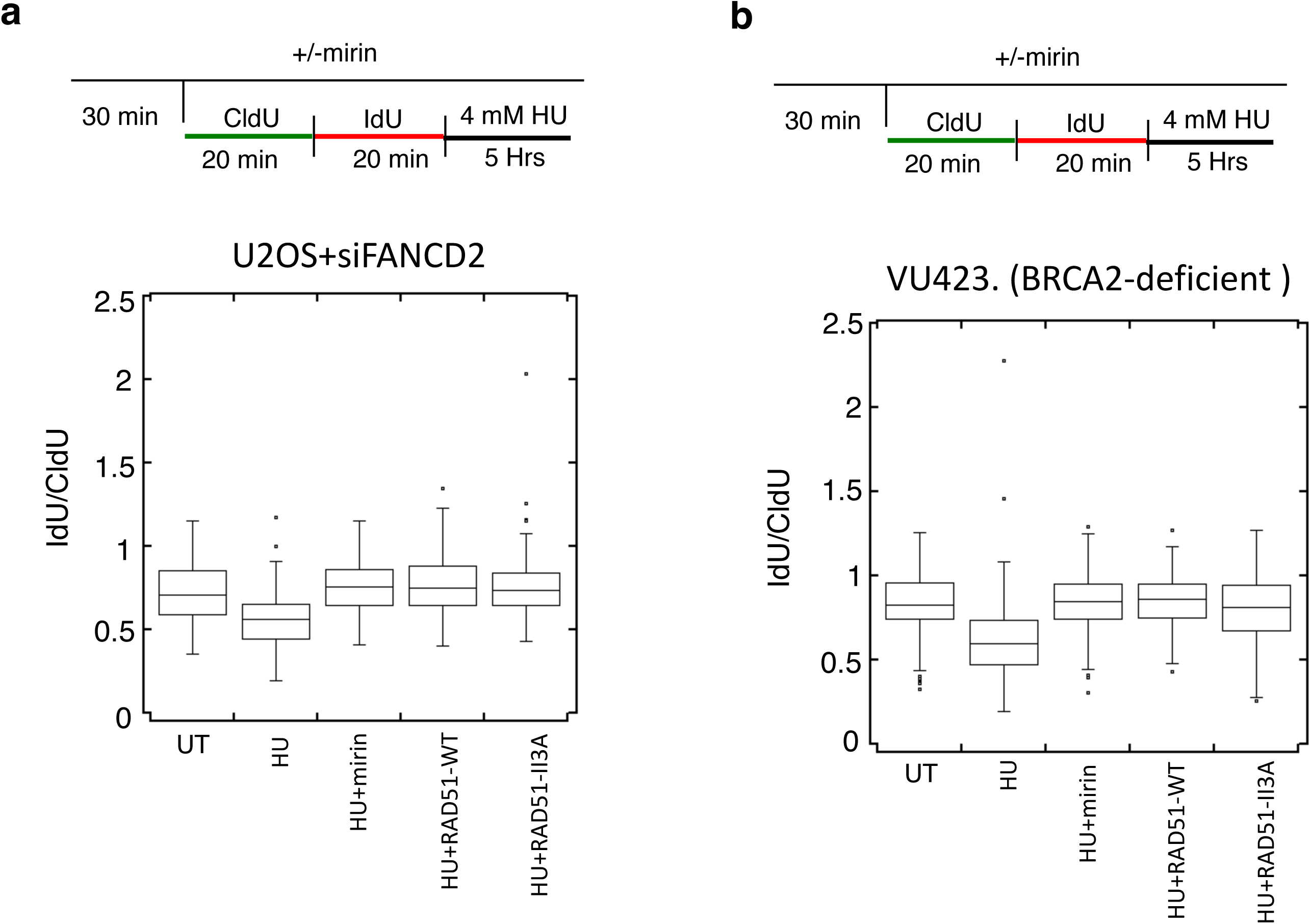
hsRAD51-II3A protects nascent strands from MRE11-dependent degradation. (a) Box plot represents the IdU/CldU ratio of tracts measured in FANCD2-depleted U2OS cells after the indicated treatment. (b) Box plot represents the IdU/CldU ratio of tracts measured in BRCA2-deficient fibroblasts (VU423) after the indicated treatment. Lines represent the median for each set of measurements. Whiskers represent 25^th^ and 75^th^ quartiles.

### hsRAD51-II3A replication tracts undergo DNA2-dependent degradation after prolonged HU treatment

MRE11-dependent degradation in human cells can be prevented by inhibiting replication fork reversal or by stabilizing RAD51 filaments on a reversed forks^7,12,14,15,17^. To distinguish between these two possibilities, we examined replication tract lengths in hsRAD51-II3A expressing cells after prolonged HU treatment. Prolonged exposure to HU (4 mM HU for 8 hours) was previously shown to result in nascent strand degradation by the nuclease DNA2 in U2OS cells without manipulations that decrease RAD51 levels or filament stability^6^. Under these conditions, DNA2-dependent degradation is dependent on RAD51-mediated fork reversal^6^. As indicated by a previous study^6^, the ability of DNA2 to degrade forks after prolonged HU treatment can be used as a readout of efficient fork reversal in U2OS cells^8,11^.

We examined fork degradation by pulsing cells with CldU followed by IdU before treatment with HU for 8 hours and measured the ratio of IdU to CldU (Figure 3a). In both siNS and hsRAD51-WT expressing cells, prolonged HU treatment resulted in a significant reduction in the CldU:IdU ratio (0.68±0.26 µm to 0.48±0.20 µm for siNS *p-*value <0.005, 0.71±0.22 µm to 0.46±0.20 µm for hsRAD51-WT *p-*value <0.005). Reducing DNA2 expression restored ratios to those observed in untreated samples. In contrast, treatment with mirin did not result in a significant change in the CldU:IdU ratio in either siNS or hsRAD51-WT expressing cells (Figure 3a, Supplemental Figure 2b). These results confirm DNA2, and not MRE11 is responsible for degradation under these conditions. Depletion of RAD51 did not result in tract degradation under any treatment condition. These findings confirm that RAD51 remodels HU stalled replication forks to provide a substrate for DNA2-mediated degradation^6^. hsRAD51-II3A expressing cells exhibited a significantly reduced CldU:IdU ratio (0.75±0.62 µm to 0.62±0.30 µm, *p-*value <0.005) that was rescued by depletion of DNA2. Treatment with mirin has no effect on the CldU:IdU ratio in hsRAD51-II3A expressing cells. We obtained an equivalent result when we measured total tract length after pulsing cells with CldU (Supplemental Figure 2c). These results confirm that, in contrast to expectation^4^, the strand exchange activity of RAD51 is not required to remodel stalled replication forks to a form that is sensitive to DNA2-mediated degradation.

**Figure 3.**
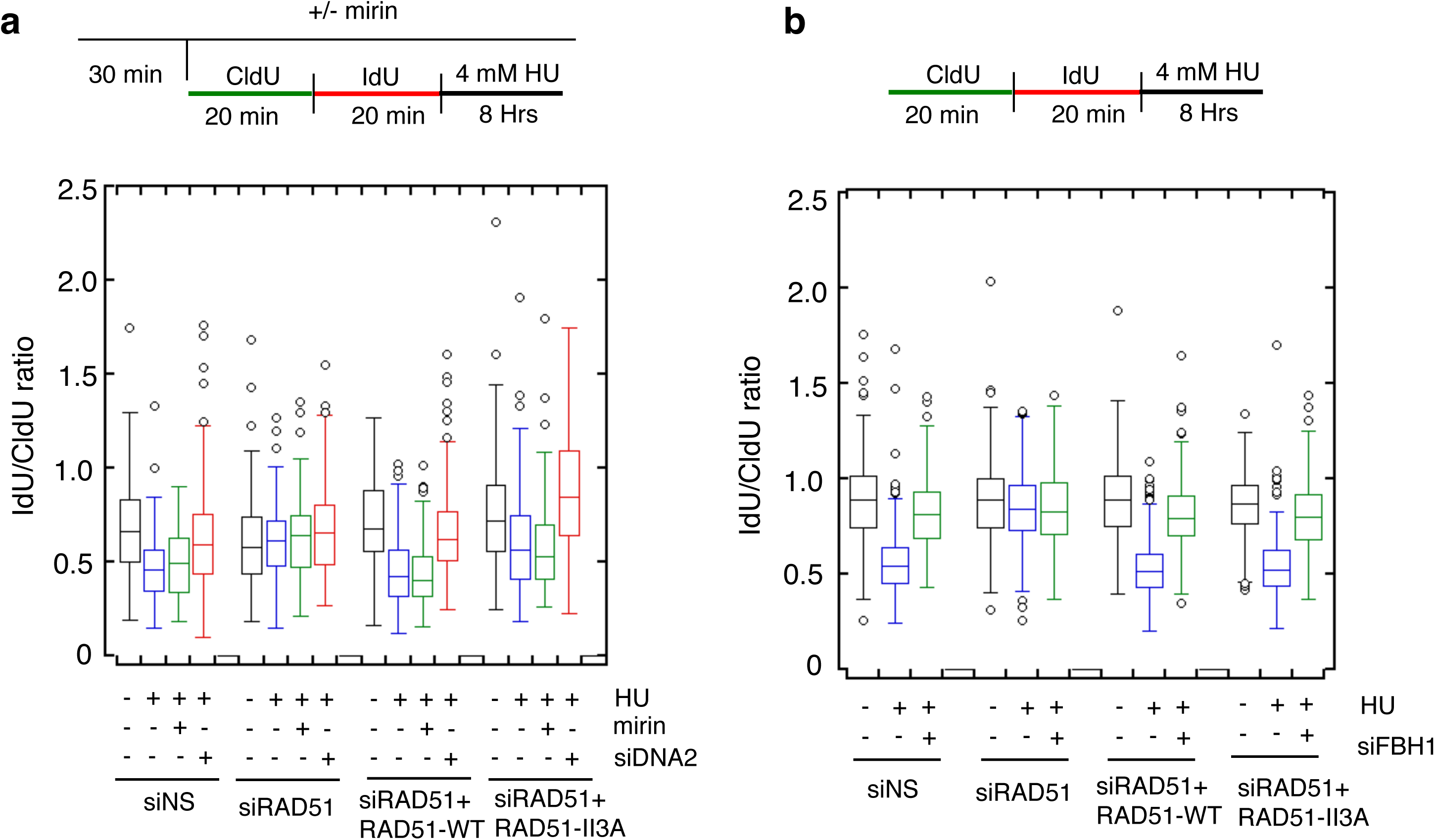
Replication tract shortening after prolonged HU treatment requires DNA2 and FBH1 activity. (a) DNA2 depletion prevents replication tract shortening after treatment with 4 mM HU for 8 hours. Box plot represents the IdU/CldU ratio of tracts measured in U2OS cells after the indicated treatment. (b) FBH1 depletion prevents replication tract shortening after treatment with 4 mM HU for 8 hours. Box plot represents the IdU/CldU ratio of tracts measured in U2OS cells after the indicated treatment. Lines represent the median for each set of measurements. Whiskers represent 25^th^ and 75^th^ quartiles.

Several proteins have been shown to catalyze fork reversal in response to hydroxyurea including the SNF2 family translocases and the helicase FBH1^15,39,40^. To determine if FBH1 functions in hsRAD51-II3A expressing cells to mediate fork reversal, we examined replication tract lengths in hsRAD51-II3A expressing cells depleted of FBH1 after prolonged HU treatment (Figure 3b, Supplemental Figure 2d). As before, treatment of siNS transfected cells with HU reduced the Cldu:IdU ratio (0.89±0.22 µm to 0.56±0.17, *p<*0.0001). Depletion of FBH1 restored tract length to those observed in untreated controls (0.82±0.17 µm) indicating FBH1 promotes DNA2-dependent degradation. As above, we did not observe degradation in siRAD51 transfected cells under any conditions. HU treatment of hsRAD51-WT (0.90±0.22 µm to 0.52±0.15, *p<*0.0001) and hsRAD51-II3A (0.86±0.17 µm to 0.54±0.15, *p<*0.0001) reduced the replication tract length and FBH1-depletion restored replication tract length in both cell lines. Thus, FBH1 promotes DNA2-dependent degradation in cells lacking RAD51 strand exchange activity, but this activity of FBH1 depends on the DNA binding activity of RAD51.

### RAD51 strand exchange activity is required for replication fork restart after prolonged HU treatment

Previous studies showed RAD51 is required for restart of replication forks stalled by HU treatment^19^. Thus, we determined if the strand exchange activity of RAD51 is required to restart stalled replication forks after treatment with HU for 5 and 8 hours (Figure 4). The siNS and hsRAD51-WT transfected cells showed significant levels of restart after both 5 and 8 hours of HU treatment; in siNS control cells 50±8 % and 39±3.9 % of replication forks restarted after 5 and 8 hours of HU treatment respectively; in hsRAD51-WT transfected cells, the corresponding numbers were 48±3.3% and 40±3.9% respectively. Depletion of RAD51 resulted in a 2-fold reduction in the frequency of restart at 5 hours (24.3±4.8% fork restart; *p*<0.005) and a 2.3-fold reduction at 8 hours (17.1±2.4% fork restart; *p*<0.005), confirming that RAD51 is required for efficient fork restart after HU treatment. hsRAD51-II3A expressing cells gave results that were intermediate between the positive and negative controls, with only a slight decrease in the efficiency of replication fork restart (44±2.6%; *p<0.05*) at 5 hours and a more severe (2.8-fold) reduction in the frequency of restart after 8 hours HU treatment (13.8±2.7% fork restart; *p<0.005)*. Thus, RAD51’s strand exchange activity is required for efficient replication restart, with a much greater requirement after 8 hours as compared to 5 hours of HU treatment. The results also raise the possibility that the strand exchange defective form of RAD51 can promote more restart than occurs when RAD51 levels are dramatically repressed. The alternative possibility is that hsRAD51-II3A has residual strand exchange activity *in vivo*, in spite of our inability to detect such activity biochemically. However, this seems unlikely because hsRAD51-II3A did not exhibit higher HR activity *in vivo* using the DR-GFP assay compared to RAD51-depleted cells (Figure 1).

**Figure 4.**
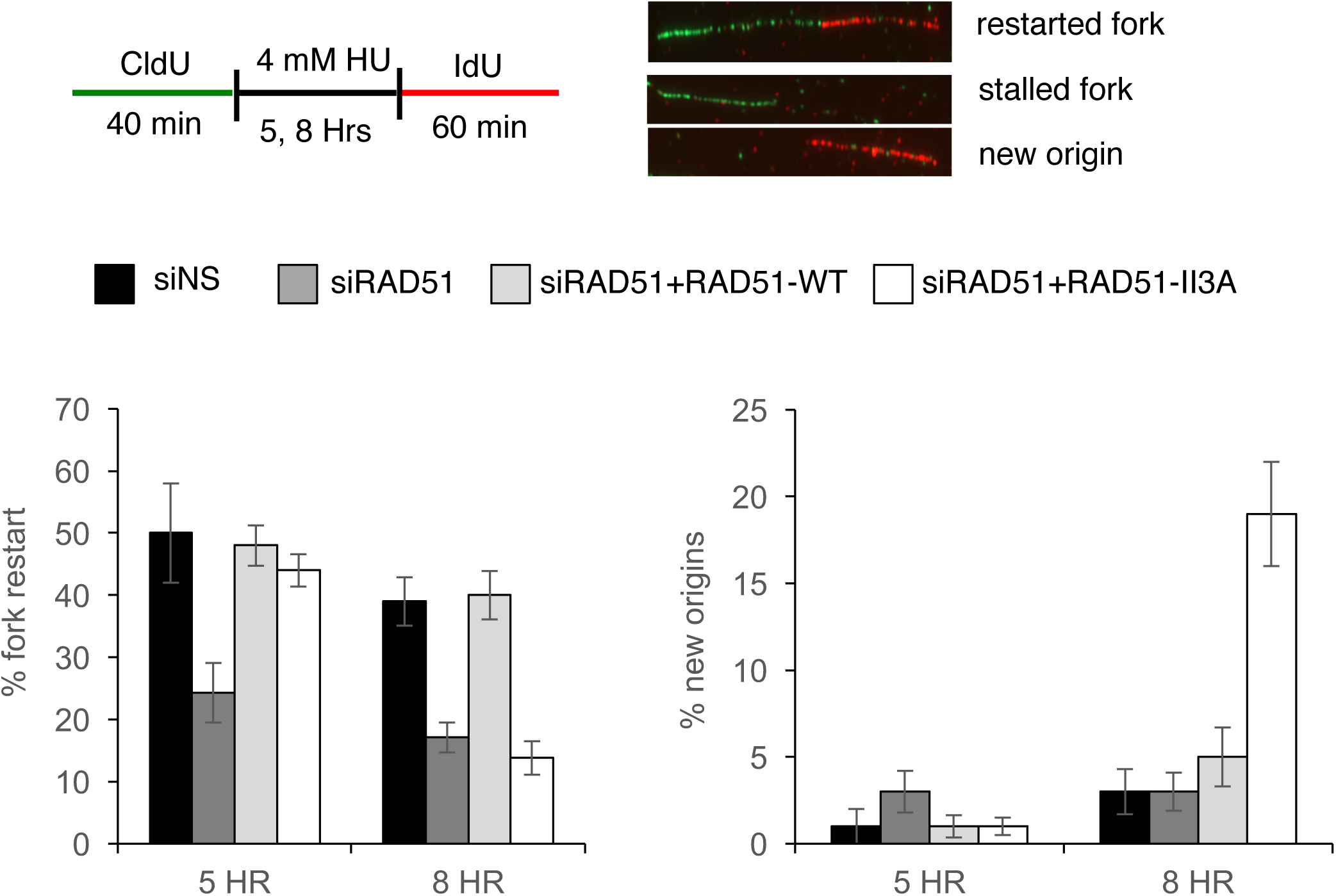
Replication restart defects in hsRAD51-II3A expressing cells after treatment with HU. Graphs depict the percentage of replication forks that restart after the indicated treatments (left) or the percentage of new origin firings (right). Error bars represent SEM. Schematic of experimental design is above the graphs and fibers that represent restarted forks (CldU+IdU), stalled fork (CldU only) and new origin (IdU only) are shown.

Next, we examined cells for new origin firing following a period of replication blockage by HU. New origin firing can be detected in the same double labeling experiments described above, by the presence of tracts containing only IdU labeling. We observed very little or no new origin firing (<5%) in the positive and negative control experiments (Figure 4). After 5 hours HU treatment, hsRAD51-II3A cells exhibited new origin firing similar to control cell lines. In contrast, hsRAD51-II3A expressing cells exhibited a 7-fold higher level of new origin firing at 8 hours as compared to siNS (19±2.7%; *p<*0.005). These data indicate that replication fork blockage by HU leads to more new origin firing in cells expressing hsRAD51-II3A, than in cells expressing hsRAD51-WT or in cells blocked for RAD51 expression.

### 53BP1 foci accumulate in hsRAD51-II3A cells after HU treatment

Replication fork blockage by HU can lead to fork collapse, a process that creates a broken DNA end that recruits DNA break proteins including 53BP1^41-45^. Replication stress-induced fork collapse and associated DNA break signaling have been shown to lead to the firing of new origins^19^. Given prior evidence for a functional association between new origin firing and fork collapse, we hypothesized that the new origin firing we observed in HU-treated hsRAD51-II3A cells is a consequence of an increased frequency of fork collapse. We therefore tested for evidence of collapsed fork accumulation specifically in S-phase cells by staining with the 53BP1, which is known to localize to DSBs, and the replication fork specific marker PCNA^46,47^. After 8 hours in HU, the siNS and hsRAD51-WT positive controls, and the siRAD51 negative control, showed a 7-fold increase in the average number of 53BP1 foci /cell (20.5±9.6 53BP1 foci/cell compared to 2.9±3.8 foci per cell prior to HU treatment; Figure 5). Expression of hsRAD51-II3A cells resulted in a significantly greater (11-fold) increase in 53BP1 foci after 8 hours HU treatment (47±0.8 53BP1 foci/cell; *p*-value<0.005) as compared to the untreated sample. In addition to being recruited at broken DNA ends formed by fork collapse, 53BP1 may form foci at DNA ends of reversed forks (i.e. the “middle toe” of the chicken foot)^44^, and is recruited to stalled forks early in the replication response^48^. As a means of determining if the observed 53BP1 foci that accumulate in RAD51-II3A expressing cells were due to fork collapse mediated by MUS81, we asked if the observed accumulation could be reduced or eliminated by siMUS81 (Figure 5, Supplemental Figure 3). Indeed, depletion of MUS81 reduced the number 53BP1 foci in siNS, siRAD51, and hsRAD51-WT expressing cells 1.8-fold (12.3±7.3 foci/cell; *p-*value<0.005). In RAD51-II3A expressing cells, MUS81 depletion resulted in a 1.9-fold decrease in the number of 53BP1 foci per nucleus (24.5±11.8 foci/cell; *p-*value<0.005) providing evidence that 53BP1 focus formation is associated with MUS81-dependent cleavage. As a control against the possibility that 53BP1 activity differs between cultures, a fraction of each culture was treated with HU for 24 hours. This highly prolonged replication arrest caused equivalent induction of 53BP1 foci and MUS81 depletion significantly reduced 53BP1 foci in all samples, as expected (Supplemental Figure 3). Together, the results suggest that hsRAD51-II3A causes more accumulation of collapsed forks following 8 hr HU treatment than occurs in cells expressing equivalent levels of hsRAD51-WT, and also more than in cells expressing very low levels of RAD51. The possible mechanistic basis for these observations is discussed below.

**Figure 5.**
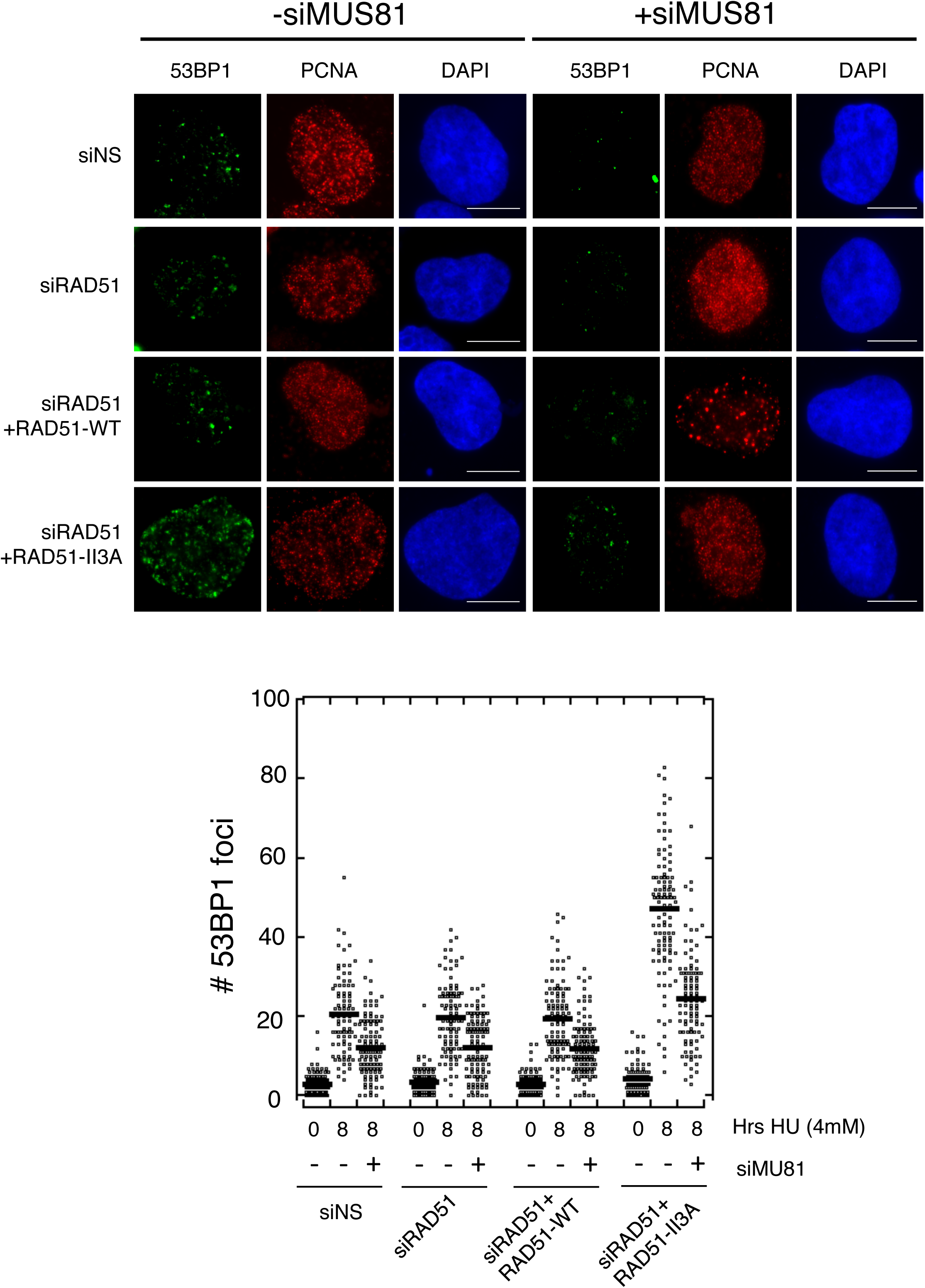
hsRAD51-II3A expressing cells accumulate 53BP1 foci after treatment with HU. Representative images depicting 53BP1 foci (green) in PCNA (red) positive cells after the indicated treatments. DNA is stained in blue. Scale bar= 10 µm. Dot plot depicts quantitation of 53BP1 foci after indicated treatments. Black line represents the mean.

## DISCUSSION

RAD51 has been implicated in several steps in the response to replication stress including fork protection, replication fork remodeling, and replication fork restart. Here, we utilized a RAD51 mutant allele that retains DNA binding activity, but is defective in strand exchange to gain mechanistic insight into the role of RAD51 at stalled replication forks. Previous studies have suggested that stabilization of RAD51 filaments is sufficient to protect from MRE11 dependent-degradation^7,12^. This study utilized the RAD51-K133R mutant that is defective for HR-mediated repair, but has robust strand exchange activity precluding the use of this mutant to definitively determine if RAD51 DNA binding activity alone is sufficient to protect from MRE11-dependent degradation^26,27^. We found that the ability of RAD51 to protect nascent strands from MRE11-mediated degradation is independent of strand exchange activity in both BRCA2- and FANCD2-deficient cell lines supporting the hypothesis that RAD51 filament formation is sufficient to protect replication forks from pathological degradation. Our results provide additional insight into the mechanism of RAD51-dependent replication fork remodeling by showing that degradation of reversed forks by DNA2 does not require RAD51’s strand exchange activity. We further show that the strand exchange activity of RAD51 is required for efficient replication restart after prolonged HU exposure.

Prolonged HU treatment of cells results in DNA2-dependent degradation that is not dependent on BRCA2-RAD51 fork protection, but depends on RAD51-dependent replication fork reversal^6,11^. We provide evidence that the strand exchange activity of RAD51 is not required for DNA2-dependent degradation of stalled forks under conditions of prolonged replication stress suggesting that DNA binding activity of RAD51 is sufficient to promote replication fork reversal. How can the DNA binding activity of RAD51 promote replication fork reversal? RAD51 interacts with polymerase a preventing the formation of ssDNA gaps at stalled forks^15^. If annealing of complementary nascent strands is important to drive fork reversal, RAD51 preventing significant ssDNA formation at the fork may be sufficient to drive fork reversal. Alternatively, DNA-bound RAD51 could promote fork reversal by recruiting other proteins that act directly to catalyze the process. RAD54^49^, FANCM^50^, HTLF^51^, and ZRANB3^52,53^, have been found to be able to reverse a model replication fork substrate *in vitro* and FBH1^39^, SMARCL1^14^, and ZRANB3^40^ have been shown to promote fork reversal *in vivo.* Here we show FBH1 depletion in RAD51-II3A expressing cells prevents DNA2-dependent degradation, providing evidence for a pathway of fork reversal that depends on both the non-enzymatic activity of RAD51 and on FBH1. Importantly we see no difference the level of DNA2 sensitivity in cells blocked for FBH1 expression when cells expressing RAD51-WT or RAD51-II3A are compared. This finding implies that the strand exchange activity of RAD51 cannot substitute for FBH1 in promoting fork reversal under these conditions. We cannot rule out the possibility that RAD51 strand exchange activity is capable of promoting fork reversal in other types of human cells, but that strand exchange-independent mechanisms involving fork remodeling proteins such as SMARCL1, ZRANB3, and FBH1 predominate in U2OS cells.

hsRAD51-II3A cells promoted significant restart after 5 hours HU treatment, but were highly defective in replication restart after longer (8 hours) treatment with HU. Together, our results lead us to a model for three distinct pathways to restart stalled replication forks, one that is RAD51-dependent, strand exchange-<u>dependent</u>; a second that is RAD51-dependent, strand exchange-<u>independent</u>; and a third that is RAD51-independent. Further, our results indicate that the RAD51-dependent, strand exchange-dependent mechanism is more predominant after 8 hours of exposure to HU as compared to 5 hours of exposure, while the converse is true for the RAD51-dependent, strand exchange-independent mechanism. These data are consistent with a previous study that identified an early, cleavage-independent restart pathway involving 53BP1 and a late, HR-dependent pathway involving BRCA1^48^. Further studies are required to determine if the role of RAD51 early in the replication stress response depends on 53BP1 or if this pathway represents the RAD51-independent pathway identified by our results. Our results are also consistent with work using an allele of *S pombe rad51* that was modelled on *S. cerevisiae* Rad51-II3A, but not biochemically characterized. That work led to the proposal that strand exchange activity coded by *S. pombe rad51*^*+*^ is dispensable for replication fork protection from Exo1, but required for efficient fork restart^54^.

Combining all the data, we propose the following model for RAD51-dependent replication fork remodeling and restart (Figure 6). At early times after fork blockage, binding of RAD51 to DNA is sufficient to protect the replisome by preventing excessive uncoupling of the replication fork; thereby preventing significant ssDNA accumulation. After fork reversal and processing of regressed arms by MRE11, EXO1, and/or DNA2, RAD51 loading onto the regressed arm prevents pathological degradation by nucleases. When the replication block is removed, reversed forks can be resolved by the action of helicases such as RECQ1, reinstating the replication fork^55^. In contrast, prolonged stalling of a replication fork results in the formation of an intermediate that requires the strand exchange activity of RAD51 for restart. One possibility is that MRE11- and DNA2-mediated resection of the “middle toe” of the reversed fork provides a single-stranded overhang that serves as a substrate for formation of a RAD51 nucleoprotein filament. In this instance, RAD51-mediated strand invasion is used to reinstate the replication fork. Alternatively, endonucleolytic cleavage of reversed fork intermediates by nucleases such as MUS81 and SLX4 may form collapsed fork structures containing single-ended DNA breaks^56,57^. These structures are expected to require RAD51-mediated strand exchange to restore functional forks^19^. Consistent with this, hsRAD51-II3A expressing cells accumulate 53BP1 foci that are dependent on MUS81 activity, indicating that a substantial fraction of the accumulated foci represents collapsed replication forks formed by MUS81 cleavage. hsRAD51-II3A expressing cells also exhibit increased origin firing. These phenotypes are associated with the accumulation of collapsed replication forks^19^. Interestingly, the collapsed fork-associated phenotypes observed in hsRAD51-II3A expressing cells are more severe than those observed in RAD51-depleted cells. This observation suggests that replication fork remodeling mediated by hsRAD51-II3A traps replication restart intermediates that cannot be resolved by RAD51 strand exchange-independent pathways. The partial dependency of 53BP1 foci on MUS81 suggests that a major fraction of the trapped intermediates are collapsed forks, but it is possible that the processing of unbroken reversed forks is also blocked by RAD51-II3A, given prior evidence that 53BP1 can localize to replication forks prior to DSB formation^44,48^.

**Figure 6.**
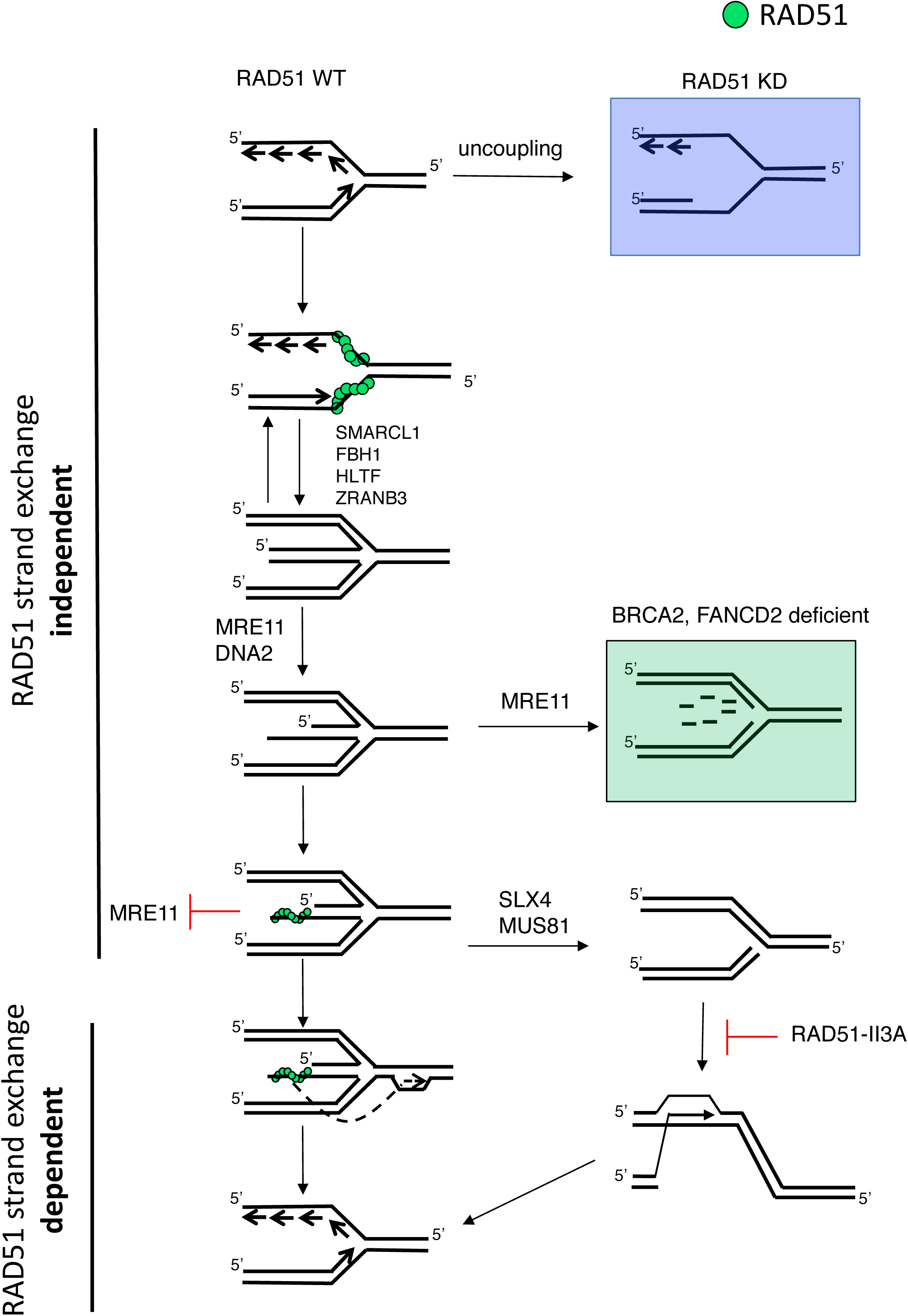
Model depicting role of RAD51 at stalled replication forks. RAD51 binds to a stalled replication fork and stabilizes it by preventing extensive ssDNA formation. Proteins such as SMARCL1 and FBH1 promote replication fork reversal resulting in the formation of the chicken foot structure. DNA2 resects the middle toe providing a substrate for RAD51 strand-exchange dependent fork restart. DNA2 also promotes degradation of nascent strands after fork reversal in cells expressing hsRAD51-WT or hsRAD51-II3A (not depicted). In RAD51 depleted cells (RAD51 KD), the replication fork contains excess ssDNA due to uncoupling (Blue Box). hsRAD51-II3A is able to prevent MRE11-degradation by binding directly to the middle toe. hsRAD51-II3A is unable to restart the fork via strand exchange activity resulting in trapped replication intermediate that often consists of a one-ended DSB formed by MUS81-dependent cleavage of a reversed fork. The one-ended DSB formed requires RAD51 strand exchange activity to repair the break and reinstate the replication fork.

Here, we demonstrate RAD51 DNA binding activity alone is sufficient for replication fork protection and reversal, but strand exchange activity is required for a significant fraction of replication fork restart. Future work will determine precisely what types of replication-associated structures require the strand exchange activity of RAD51. It will also be of interest to determine if strand exchange activity of RAD51 has additional roles at replication forks under conditions which require repair of a physical lesion (e.g. interstrand crosslinks), or conditions that only result in a moderate reduction in replication fork speed (e.g. UV-light induced damage)^10,18^.

## DATA AVAILABILITY

All data supporting the findings of this study are available from the corresponding authors upon request.

## ACKNOWLEDGEMENTS

This work is funded by the National Institutes of Health grants GM50936 to DKB, and CA205518-01 to DKB. We thank Alessandro Vindigni for helpful comments on a draft of the manuscript.

## ONLINE MATERIAL AND METHODS

### Expression and purification of hsRAD51-WT and hsRAD51-II3A mutant

The open reading frames of hsRAD51-WT and hsRAD51-II3A mutant with a C-terminal His-6 tag were cloned into pET21d (Novagen). The proteins were overexpressed in *E. coli* Rosetta(DE3) plysS cells by induction using 0.5 mM IPTG. The expression and purification were as detailed previously for protein yeast Dmc1^**58**^.

### Binding assay

The binding of hsRAD51 to ssDNA was assayed by the fluorescence polarization method as described previously^**32**^ with the following modifications. An 84-mer ssDNA conjugated with Alexa Flour-488 at the 5’ end (sequence: 5’-GGTAGCGGTTGGGTGAGTGGTGGGGAGGGTCGGGAGGTGGCGTAGAAACATGATAGGAATGTGAATGAATGAAGTACAAGTAAA-3’; synthesized by Integrated DNA Technologies) was used at 200 nM nucleotides (2.4 nM). The binding reactions were performed at 37°C for 30 minutes in buffer B (25 mM Tris-HCl (pH 7.8), 1 mM MgCl_2_, 1 mM ATP, 1 mM DTT, 50 mM NaCl_2_, 50 µM CaCl_2_, and 100 µg/ml BSA). The fluorescence polarization (in mP units) was measured using a Tecan Infinite F200 PRO plate reader. All binding conditions were performed in triplicate, and the mean values were plotted with standard deviation. Buffer and ssDNA had no effect on fluorescence polarization in the absence of added protein (data not shown). The first data point on the graph contains 10 nM protein.

### D-loop assay

The assay was performed essentially as described previously^58^. Reactions were carried out in 25 mM Tris-HCl (pH 7.8); 1 mM MgCl_2_, 1 mM ATP, 1 mM DTT, 50 µM CaCl_2_, and 100 µg/ml BSA; ssDNA (90 mer sequence 5’TACGAATGCACACGGTGTGGTGGGCCCAGGTATTGTTAGCGGTTTGAAGCAGGCGGCAGAAGAAGTAACAAAGGAACCTAGAGGCCTTTT) was used at 3.6 µM nucleotide or 40 nM); negative supercoiled plasmid was pRS306 at 5 nM (22 µM bp).

### Cell culture

U2OS DR-GFP cells were grown in DMEM (Gibco) supplemented with 10% Fetal Bovine Serum. The authenticity of the cell line was validated previously by short tandem repeat profiling at the Genetic Resources Core Facility at John Hopkins School of Medicine (Baltimore, MD)^59^. Cell line was monitored monthly for mycoplasma contamination.

### Expression of RAD51 in U2OS cells

WT RAD51 or RAD51 cDNA containing mutations in the secondary binding site (R130A, K303A, R310A) was cloned into pcDNA 3.1 (Invitrogen) using Gibson assembly per manufacturer’s instructions (New England Biolabs). U2OS cells were transfected with pcDNA3.1, hsRAD51-WT (pNRB707), or hsRAD51-II3A (pNRB708) expression plasmids using Lipofectamine 3000 (Invitrogen) as per manufacturer’s instructions. After 24 hours, cells were transfected with RAD51 siRNAs targeting the 3’UTR. At 48-hours post transfection, cells were collected and analyzed for the various assays.

### siRNA sequences

siRNAs were transfected using Lipofetamine RNAiMAX as per manufacturer’s instructions (Invitrogen). The All-Star negative control (siNS) siRNA was used as a control (Qiagen). The following siRNA sequences were used in this study.

siRAD51 5’ GACUGCCAGGAUAAAGCUU was used in a previous study^60^.

siDNA2 5’ CAGUAUCUCCUCUAGCUAG was used in a previous study^6^.

siFANCD2 5’ CAGAGUUUGCUUCACUCUCUA was used in a previous study^61^

siMUS81 5’ CAGCCCUGGUGGAUCGAUA was used in a previous study^62^.

siFBH1 5’ GGAUGUUUGCAAGAGAGUCAGGAAA

### Nascent DNA fiber assay

Cells were pulsed with CldU (50 µM), or CldU (50 µM) followed by IdU (150 µM) were treated with HU (4mM) for the indicated times. Tract lengths were measured using Image J. To measure replication restart, cells were pulsed with CldU (50 µM) before treatment with 4mM HU for the indicated times. HU was removed and cells were pulsed with IdU (50 µM). Mirin (50 µM) was added 30 minutes prior to the pulse with CldU and was present throughout the experiment. The nascent DNA fiber assay was performed as previously described^33^. At least 150 replication tracts were measured for each condition. The plots represent pooled data from at least two independent experiments. Statistical significance was determined using Mann-Whitney U test.

### RAD51 and 53BP1 focus formation

48 hours after transfection with siRNAs, cells were treated with 4 mM HU for the indicated times. For RAD51 focus formation, cells were treated with 6 Gy using a maxitron x-ray generator. Cells were fixed and stained as previously described^33^. Dot plots represent 50 nuclei combined from two independent experiments. As with the pooled data, expression of RAD51-WT and RAD51-II3A exhibited significantly more RAD51 foci in the individual, separate experiments when compared to siRAD51 (*p*<0.05), but did not significantly differ from irradiated siNS cells (*p*>0.05). Antibodies used in this study are as followed: RAD51 is a rabbit polyclonal antibody against purified human RAD51 (1:1000, Pacific Immunology). 53BP1 (1:1000, NB100-304) was from Novus Biologicals and PCNA (1:1000, IG7) was from Abnova. Dot plots represent combined data from two independent experiments. Statistical significance was determined by the Wilcoxon Rank Sum Test.

### Western blotting

Western blotting was done as previously described^59^. Anti-DNA2 (1:500; ab96488), Anti-MUS81 (1:1000, ab14387), and anti-FBH1 (1:100, ab58881) were from Abcam. Anti-FANCD2 (1:1000, NB100-182) was from Novus Biologicals. Proteins were detected using a C-DIGIT blot scanner (Licor).

### DR-GFP assay

U2OS cells containing the DR-GFP construct stably integrated into the genome were transfected with a plasmid expressing I-SceI (pBAS) or an empty vector (pCAGG) after the indicated treatments^37^. After 48 hours, cells were collected and the percentage of cells expressing GFP was determined by flow cytometry (LSR II, BD Biosciences). Statistical significance was determined using a two-tailed t-test.

---

**Supplemental Figure 1.**
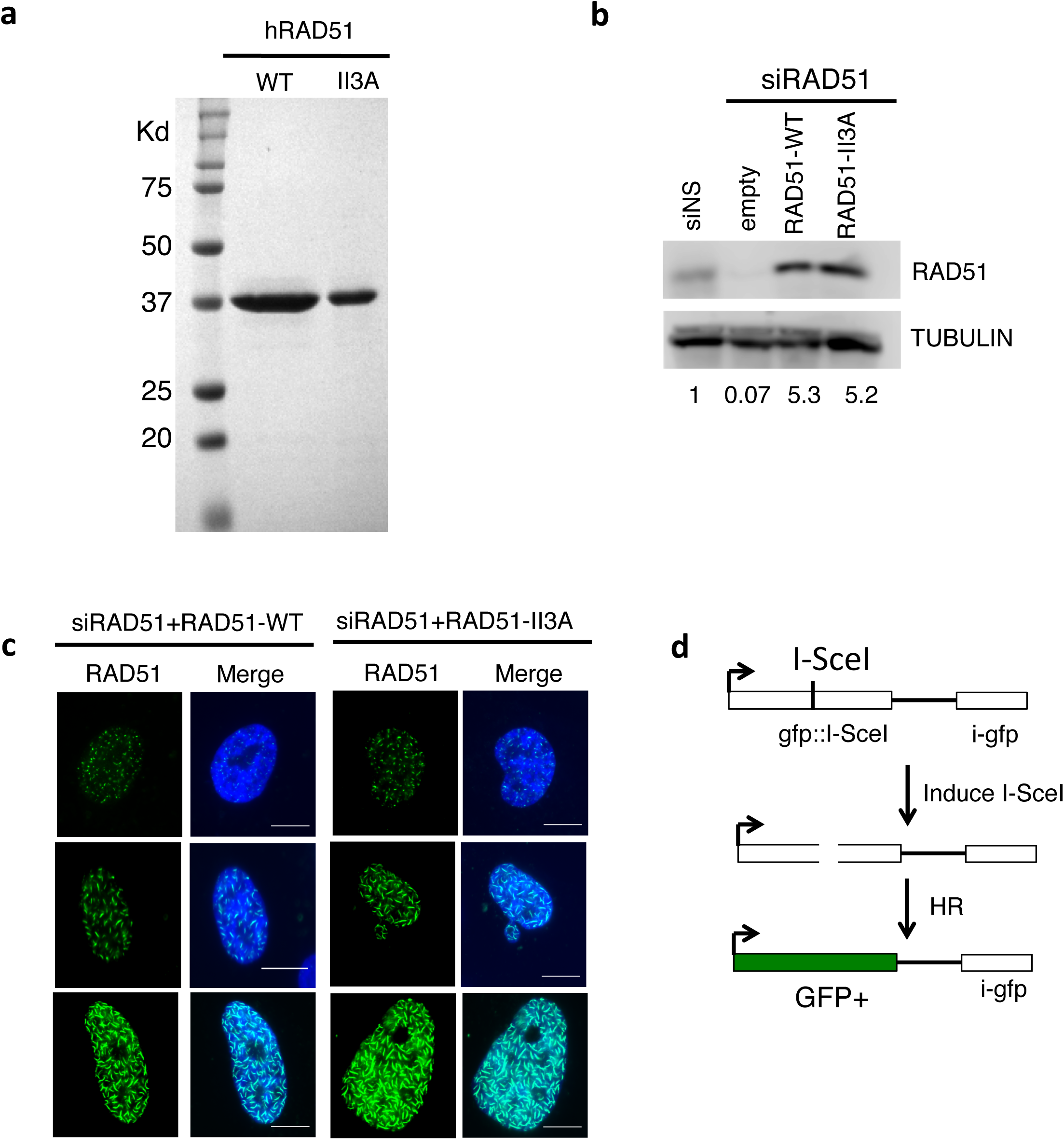
Cells over-expressing hsRAD51-WT or hsRAD51-II3A. (a) Purified hRad51-WT (25 µg) and hRad51-II3A mutant (10 µg) were analyzed on a 12% SDS-PAGE and the proteins were stained with Coomassie brilliant blue R250. (b) Western blot depicting levels of RAD51 after indicated treatments. TUBULIN was used as a loading control. The level of RAD51 protein levels (normalized to TUBULIN) relative to the siNS control are indicated below the blot. (c) Representative images of RAD51 fibers (non-damage associated complexes) in a small subpopulation of cells transfected with hsRAD51-WT or hsRAD51-II3A expression constructs Green-RAD51. Blue-DNA. Scale bar = 10 µm. (d) Depiction of the DR-GFP assay to measure homologous recombination in cells. The GFP coding sequence is disrupted by an I-SceI nuclease site and contains an internal GFP fragment downstream. Repair of the I-SceI induced DSB by HR restores the GFP coding sequence. Thus, HR efficiency is measured by determining the percentage of cells expressing GFP.

**Supplemental Figure 2.**
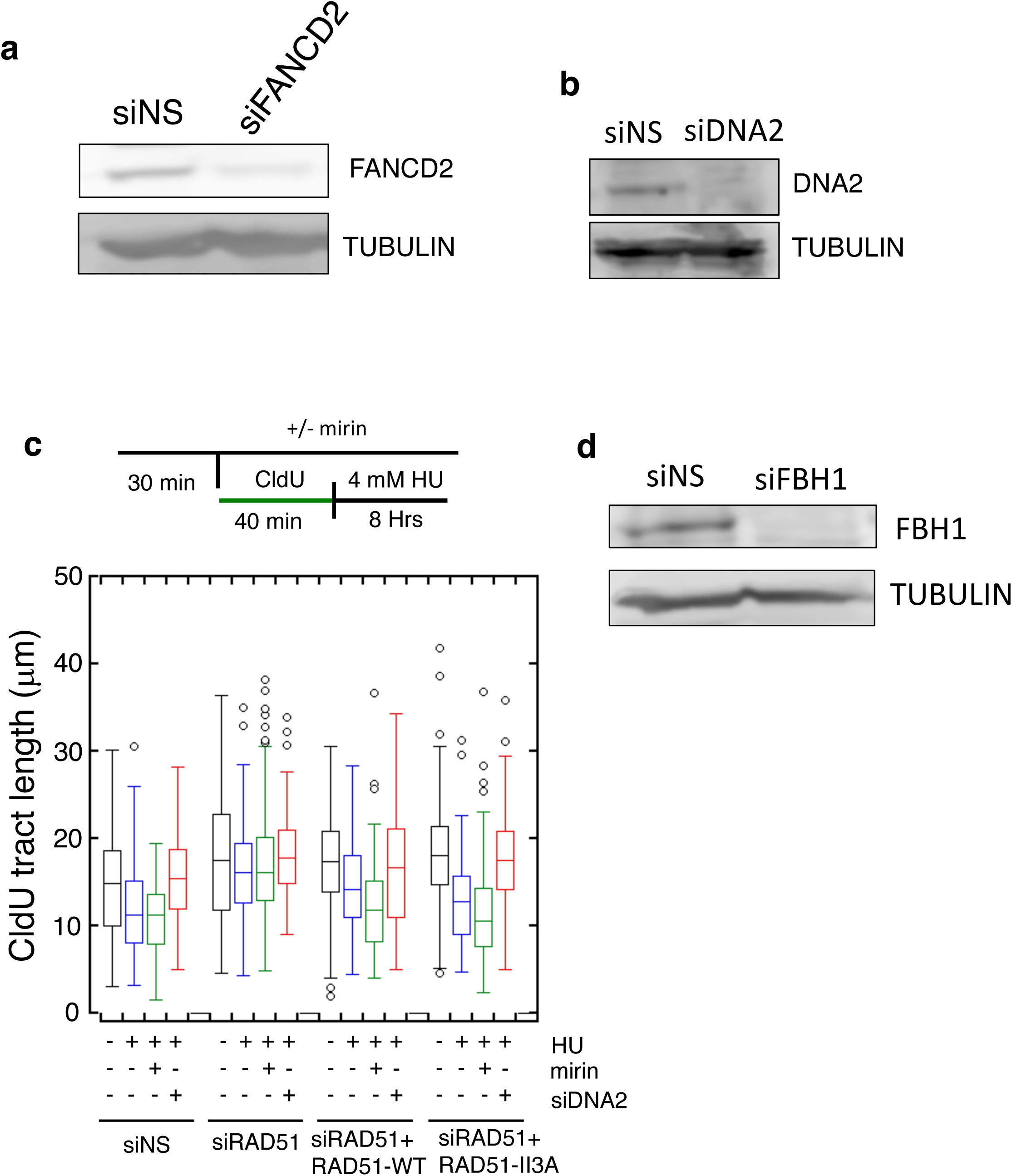
Degradation of replication tracts in RAD51-II3A is dependent on siDNA2. (a) Western depicting level of FANCD2 after indicated treatments. TUBULIN was used as a loading control. (b) Western depicting level of DNA2 after the indicated treatments. TUBULIN was used as a loading control. (c) Box plot represents the CldU tract length after the indicated treatment. Lines represent the median for each set of measurements. Whiskers represent 25^th^ and 75^th^ quartiles. Schematic of experimental design is depicted above the graphs. (d) Western depicting level of FBH1 after the indicated treatments. TUBULIN was used as a loading control.

**Supplemental Figure 3.**
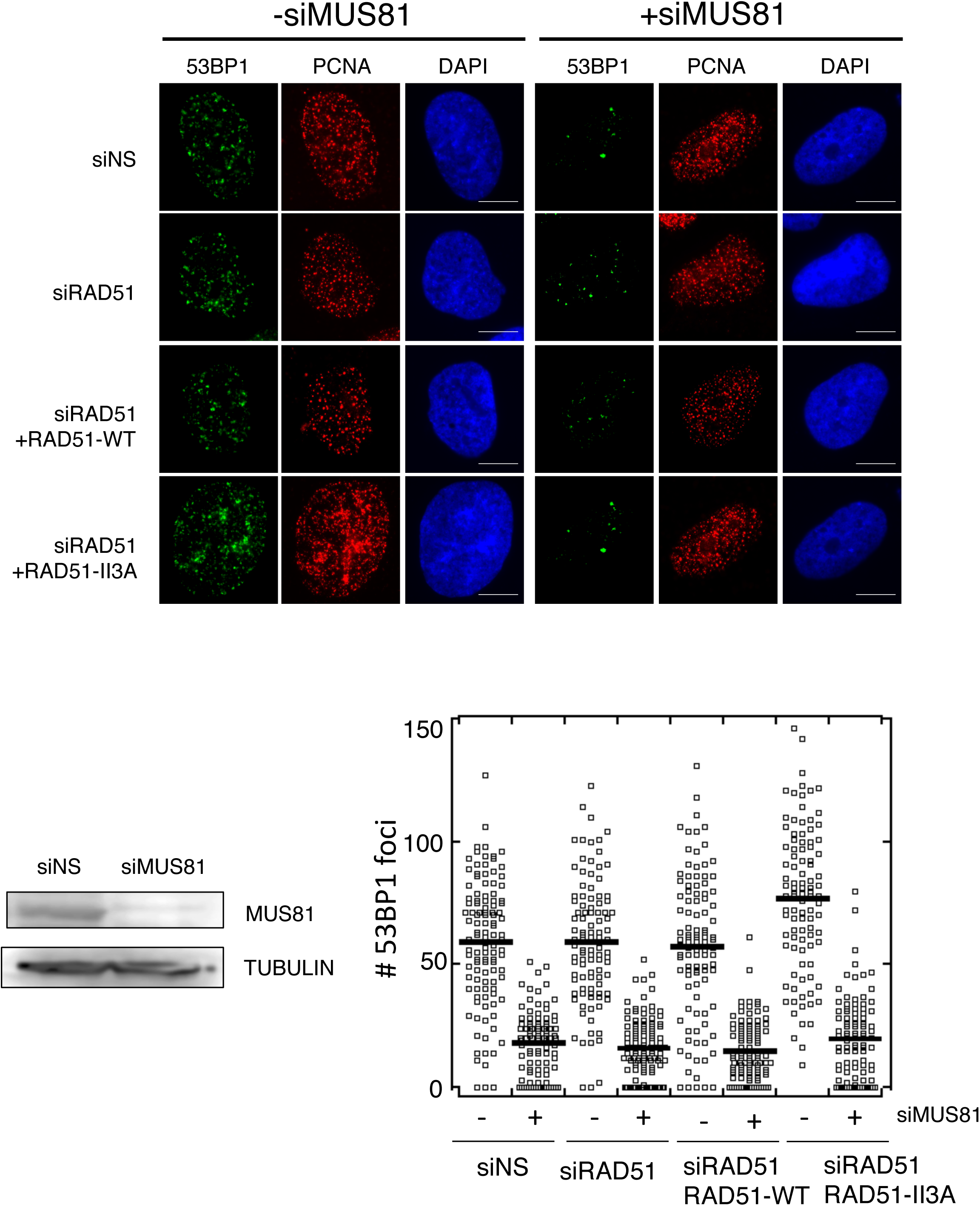
53BP1 accumulates in cells after 24 hours of HU treatment. Representative images depicting 53BP1 foci (green) in PCNA (red) positive cells after treatment with 4mM HU for 24 hours. DNA is stained in blue. Scale bar= 10 µm. Dot plot depicts quantitation of 53BP1 foci in the indicated cell lines. Line represents the mean.

